# Mitochondrial acid-sensing ion channel 1a deficiency induces mitochondrial dysfunction in pulmonary arterial smooth muscle cells

**DOI:** 10.1101/2025.09.19.677483

**Authors:** Megan N. Tuineau, Lindsay M. Herbert, Heaven E. Medina, Jay S. Naik, Thomas C. Resta, Nikki L. Jernigan

## Abstract

Pulmonary hypertension (PH) is a progressive vascular disease driven by pulmonary arterial remodeling, characterized by cellular hyperproliferation, resistance to apoptosis, and phenotypic plasticity. Our laboratory has shown that the proton-gated cation channel, acid-sensing ion channel 1a (ASIC1a), is essential for the development of chronic hypoxia (CH)-induced PH in rodents. Importantly, ASIC1a activation occurs without changes in total ASIC1a levels but reflects a hypoxia-dependent redistribution to the plasma membrane in pulmonary arterial smooth muscle cells (PASMCs). In neurons, mitochondrial-localized ASIC1a (mtASIC1a) contributes to oxidative stress-induced mitochondrial membrane potential (ΔΨm) depolarization and apoptosis. Although mtASIC1a has not been described in vascular cells, its role in PASMCs may be relevant to mitochondrial dysfunction and apoptosis resistance in PH. We hypothesize that mtASIC1a is a crucial regulator of PASMC mitochondrial homeostasis, and its loss following CH promotes mitochondrial dysfunction and apoptosis resistance. Consistent with this, mtASIC1a localization was decreased in PASMCs and intrapulmonary arteries from CH rats compared to controls. Functionally, PASMCs from CH rats or *Asic1a* knockout mice exhibited ΔΨm hyperpolarization, elevated mitochondrial Ca²⁺ and superoxide, impaired mitophagy, and reduced cleaved caspase-3. Transmission electron microscopy revealed mitochondrial morphological changes, including increased size and circularity, decreased aspect ratio, and reduced mitochondrial number per cell, while fusion/fission proteins remained largely unchanged. Lentiviral restoration of mtASIC1a prevented ΔΨm hyperpolarization and restored caspase-3 cleavage. These findings identify mtASIC1a as a novel regulator of mitochondrial function in PASMCs, where its loss following CH promotes ΔΨm hyperpolarization, impaired mitophagy, and resistance to apoptosis.

**New & Noteworthy:** This study identifies mitochondrial acid-sensing ion channel 1a (mtASIC1a) as a novel regulator of mitochondrial homeostasis in pulmonary arterial smooth muscle cells (PASMCs). Critically, mtASIC1a deficiency in PASMCs following *in vivo* chronic hypoxia or genetic deletion promotes mitochondrial membrane potential (ΔΨm) hyperpolarization, Ca²⁺ and O_2_^-^ accumulation, impaired mitophagy, and caspase inhibition. Restoring mtASIC1a by lentiviral transduction prevents ΔΨm hyperpolarization and restores caspase cleavage, highlighting its importance in mitochondrial signaling and hypoxic pulmonary hypertension pathophysiology.

## 1 Introduction

Chronic lung diseases or lifestyle factors such as chronic obstructive pulmonary disease, interstitial lung disease, sleep apnea, smoking, and high-altitude residence frequently cause prolonged alveolar hypoxia. Unfortunately, these patients often develop comorbidities, including pulmonary hypertension (PH), a condition with increasing prevalence and incidence worldwide. If left untreated, PH leads to right ventricular hypertrophy and ultimately right heart failure and mortality. Notably, patients with hypoxic PH, classified as Group III PH by the World Health Organization, have the worst long-term survival among all groups (1,2).

In PH, increased pulmonary vascular resistance primarily results from sustained constriction and remodeling of pulmonary arteries. Notably, this remodeling is characterized by cellular hyperproliferation, resistance to apoptosis, and phenotypic transdifferentiation (3–8). Although these changes arise from a complex interplay of factors, mitochondrial dysfunction, including impaired bioenergetics and oxidative stress, has emerged as a central pathogenic feature involved in these processes (9–12). Despite this central role, the upstream mechanisms responsible for the mitochondrial dysfunction observed in PH remain poorly defined.

Our prior work demonstrates that acid-sensing ion channel 1a (ASIC1a), a proton-gated, voltage-insensitive cation channel, is required for the development of chronic hypoxia (CH)-induced PH (13,14). In pulmonary arterial smooth muscle cells (PASMCs), CH enhances ASIC1a-dependent Na^+^ and Ca^2+^ influx, which is linked with plasma membrane depolarization, cell proliferation, migration, vasoconstriction, and vascular remodeling (13–17). These effects occur without changes in total ASIC1a protein levels; instead, its localization to the plasma membrane is increased (18). While the functional implications of this altered cellular localization of ASIC1a are not fully understood, evidence suggests that intracellular ASIC1a plays a distinct physiological role.

Recent studies suggest that a subpopulation of ASIC1a localizes to mitochondria, where it regulates mitochondrial-dependent cell death induced by oxidative stress in neurons (19,20). Although mitochondrial ASIC1a (mtASIC1a) has not yet been described in vascular cells, it represents a plausible intracellular pool depleted by CH-induced plasma membrane recruitment of ASIC1a. This possibility aligns with the growing recognition that mitochondrial ion channels are critical regulators of mitochondrial membrane potential (ΔΨm), mitochondrial ionic homeostasis, and apoptosis – processes intimately tied to pulmonary vascular dysfunction.

In this study, we aimed to investigate whether mtASIC1a is functionally relevant in PASMCs and whether the redistribution of ASIC1a following CH affects mitochondrial function. We hypothesize that ASIC1a is a functional mitochondrial ion channel, and its localization is decreased following CH, causing mitochondrial dysfunction and apoptosis resistance. To test this, we examined PASMCs from *Asic1a* knockout mice and from rats exposed to CH. In addition, we assessed the effects of mtASIC1a restoration using lentiviral transduction of mitochondrial-targeted ASIC1a.

## 2 Materials and Methods Materials and Methods

### 2.1 Ethical Approval

All experimental protocols were reviewed and approved by the Institutional Animal Care and Use Committee of the University of New Mexico School of Medicine (Albuquerque, NM; Protocol 25-201645-HSC) and conducted in accordance with the National Institutes of Health guidelines for animal care and use. Animals were anesthetized with an overdose of pentobarbital sodium (200 mg/kg, i.p.) and euthanized by exsanguination following the loss of consciousness.

### 2.2 Animals

Experiments were performed in either adult male Wistar rats (200–250 g, Envigo) or *Asic1a* knockout mice [B6.129-Asic1*^tm1Wsh^*/J; Jackson Laboratory Stock #013733] on a C57BL/6 background. Age-matched C57BL/6J mice (14–20 weeks; Jackson Laboratory Stock #000664) served as wild-type controls. Genotyping of homozygous and heterozygous mice was confirmed by PCR and agarose gel electrophoresis, as previously described (17). Animals were housed 1–5 per cage in a specific pathogen-free facility on a 12:12-h light-dark cycle with *ad libitum* access to standard chow (Teklad 2920X, Envigo) and water. Animals were randomly allocated to experimental groups, and when feasible, investigators were blinded to treatment assignments.

### 2.3 Model of Chronic Hypoxia-induced Pulmonary Hypertension

Animals were randomly assigned to experimental groups, and animals designated to CH exposure were housed in a clear plexiglass hypobaric chamber (∼0.5 m³) maintained at 380 mmHg for 4 weeks. The chamber was partially evacuated with a vacuum pump with continuous airflow (30 L/min). Bedding, chow, and water were replenished three times per week when the chamber was briefly opened. Age-matched control animals were maintained at ambient barometric pressure (∼630 mmHg, Albuquerque, NM). We have previously demonstrated that this model recapitulates key features of PH, including elevated right ventricular systolic pressure, right ventricular hypertrophy, enhanced pulmonary arterial vasoconstriction, and pulmonary arterial remodeling (21).

### 2.4 Isolation of Intrapulmonary Arteries and Primary Arterial Smooth Muscle Cell Culture

In anesthetized animals, the heart and lungs were excised via midline thoracotomy. Intrapulmonary arteries (IPAs; ∼ second to fifth order) were dissected free of lung parenchyma and either snap-frozen in liquid nitrogen for Western blotting or enzymatically digested for primary PASMC culture.

#### 2.4.1 Rat PASMC primary culture

Isolated arteries from control and CH rats were incubated in Hanks’ balanced salt solution (HBSS; Lonza CAT#10-547F) containing CaCl2 (20 μM), papain (26 U/ml), type I collagenase (1,450 U/ml), dithiothreitol (1 mg/ml), and bovine serum albumin (2 mg/ml) at 37°C for 30 min. Following trituration, the cell suspension was plated in Ham’s F-12 medium (Gibco) supplemented with 5% fetal bovine serum (Gibco) and 1% penicillin-streptomycin (Gibco) and transiently cultured for 3-4 days. Before all experiments, morphological appearance was assessed using phase contrast microscopy to determine cellular purity. We have previously reported expression for SM22a, smooth muscle actin, and smooth muscle myosin heavy chain in cultured PASMCs using immunofluorescence (18,22).

### 2.4.2 Mouse PASMC primary culture

Isolated arteries were incubated in HBSS containing CaCl_2_ (20 μM), papain (9.5 U/ml), type I collagenase (1,750 U/ml), dithiothreitol (1 mg/ml), and bovine serum albumin (2 mg/ml) at 37°C for 30 min. Following digestion, single smooth muscle cells were dispersed by gentle trituration with a fire-polished pipette in Ca^2+^-free HBSS. The cell suspension was plated in SmBM Basal Medium (Lonza) supplemented with SmGM-2 SingleQuots (Lonza) and 1% penicillin-streptomycin (Gibco). For these experiments, cultures were passaged twice to generate enough cells. Additionally, due to the time required to culture mouse primary PASMCs, we did not generate cultures from PH mice.

### 2.4.3 Lentiviral Transduction

For rescue experiments, a lentiviral vector carrying the sequence for human ASIC1a with a mitochondria-targeting sequence (EGFP-mtASIC1a; pLV-EGFP-CMV>MITO/hASIC1a) was generated and packaged by VectorBuilder. Freshly suspended PASMCs were plated in the appropriate serum-free media containing EGFP-mtASIC1a (∼1×10^4^ transducing units/μL), polybrene (1.5 mg/mL; VectorBuilder), and 1% penicillin-streptomycin and transduced for 24 h before medium replacement. For experiments, effective transduction was identified by EGFP fluorescence, and the experimental results were compared with those of PASMCs transduced with the control vector (EGFP-empty) supplied by VectorBuilder.

### 2.5 Cellular Fractionation and Western Blot

Cultured PASMCs or IPAs from animals were homogenized in Tris-HCl homogenization buffer [pH adjusted to 7.4 with NaOH containing 255 mM sucrose, 10 mM Tris-HCl, 2 mM EDTA, and 1X Halt Protease Inhibitor Cocktail (Thermo Fisher Scientific)] and centrifuged at 10,000 g for 10 min at 4 °C to remove insoluble debris. Protein concentrations were determined using a Qubit Protein Assay Kit (Life Technologies). Samples were heat-denatured in 1X sample buffer (60 mM Tris-HCl, pH 6.8, 25% glycerol, 2% SDS, 0.1% bromophenol blue, 5% β-mercaptoethanol), separated by SDS-PAGE, and transferred to polyvinylidene difluoride membranes. Membranes were blocked with 5% non-fat dry milk for 1 h, then incubated with primary antibodies overnight at 4 °C and with secondary antibodies at room temperature for 1 h (Table 1). Protein detection was performed by chemiluminescence (Pierce). Densitometry was performed using ImageJ, with normalization to total protein per lane by Coomassie Brilliant Blue (CBB) staining.

**Table 1.** List of primary and secondary antibodies used for immunofluorescence and western blot analysis.

| Antibody | Company | Cat # | RRID | Host; Clone | Dilution | Validation |
| --- | --- | --- | --- | --- | --- | --- |
| anti-cleaved caspase-3 | Proteintech | 25128-1-AP | AB_3073913 | rabbit; poly | 1:100 | Induction with staurosporine by manufacturer |
| Anti-LC3B | Abcam | ab192890 | AB_2827794 | rabbit; mono | 1:1,000 | knockout by manufacturer |
| anti-Parkin | Abcam | ab77924 | AB_1566559 | mouse; mono | 1:2,000 | knockout by manufacturer |
| anti-MFN1 | Proteintech | 13798-1-AP | AB_2266318 | rabbit; poly | 1:1,000 | knockout (64) |
| anti-MFN2 | Proteintech | 12186-1-AP | AB_2266320 | rabbit; poly | 1:1,000 | knockout (64) |
| anti-DRP1 | Proteintech | 12957-1-AP | AB_2093525 | rabbit; poly | 1:500 | siRNA knockdown (65) |
| anti-FIS1 | Proteintech | 10956-1-AP | AB_2102532 | rabbit; poly | 1:500 | siRNA knockdown (66) |
| anti-ASIC1a | Sigma Aldrich | AB5674P | AB_91972 | rabbit; poly | 1:500 (WB), 1:100 (IF) | siRNA knockdown (67) |
| anti-COXIV | Thermo Fisher Scientific | A21348 | AB_2535839 | rabbit; poly | 1:500 | siRNA knockdown (68) |
| anti-TOMM20 | Proteintech | CL594-11802 | AB_2919775 | rabbit;poly | 1:100 | siRNA knockdown by manufacturer |
| anti-PINK1 | Proteintech | 23274-1-AP | AB_2879244 | rabbit; poly | 1:500 | siRNA knockdown by manufacturer |
| anti-Rabbit IgG (H+L)-HRP Conjugate | Bio-Rad | 1721019 | AB_11125143 | goat; poly | 1:3,000 |  |
| anti-Mouse IgG (H+L)-HRP Conjugate | Bio-Rad | 1721001 | AB_11125936 | goat; poly | 1:3,000 |  |

For some experiments, mitochondrial-enriched fractions were isolated from IPAs or cultured PASMCs. Samples were homogenized in Tris-HCl homogenization buffer and centrifuged at 600 g for 5 min. The supernatants were removed and centrifuged at 10,000 g for 20 minutes.

Mitochondrial-enriched pellets were resuspended in homogenization buffer before use for western blot analysis.

### 2.6 Mitochondrial Membrane Potential, Ca^2+^, and superoxide (O_2_^-^)

To assess mitochondrial function, we examined ΔΨm, mitochondrial Ca^2+^ levels ([Ca^2+^]*_mito_*), and mitochondrial O_2_^-^ levels ([O_2_^-^]*_mito_*) using fluorescent-based indicators in cultured PASMCs. Before the experiments, PASMCs were cultured in serum-free medium for 24 hours to reduce the confounding effects of serum-derived growth factors and synchronize cells in a quiescent (G_0_) phase. PASMCs were incubated at 37 °C in HEPES-based physiological saline solution (PSS) [(in mM) 130 NaCl, 4 KC1, 1.2 MgSO_4_, 4 NaHCO_3_, 10 HEPES, 1.18 KH_2_PO_4_, 1.8 CaCl_2_, 6 glucose] containing either:

Tetramethylrhodamine ethyl ester (TMRE, Thermo Fisher Scientific) at 1 μM for 30 minutes. PASMCs were washed before live-cell imaging of ΔΨm. In some experiments, TMRE fluorescence was measured in PASMCs superfused with PSS (5 mL/min, 37°C) before the addition of the protonophore 4-trifluoromethoxyphenylhydrazone (FCCP; 20 μM, Cayman Chemica). Samples were excited using at 540 ± 30 nm using an IonOptix Hyperswitch light source (IonOptix LLC), and 565 ± 20-nm emissions were collected with a photomultiplier tube. Otherwise, TMRE (excitation: 552 nm; emission 574 nm) was detected using a confocal microscopy system (Leica TCS SP5).

Rhod-2 AM (Thermo Fisher Scientific)at 2.5 μM with 0.05% pluronic acid for 30 minutes. PASMCs were washed and incubated for an additional 30 min in dye-free HEPES-based PSS to allow for ester hydrolysis before live-cell imaging of [Ca^2+^]*_mito_* (excitation: 552 nm; emission: 581 nm) using a confocal microscopy system (Leica TCS SP5).

MitoSOX (Thermo Fisher Scientific)at 1 μM with 0.05% pluronic acid for 10 minutes. PASMCs were washed and fixed with 2% paraformaldehyde for 15 min, washed, mounted, and relative [O_2_^-^]*_mito_* was imaged (excitation: 510 nm; emission: 580 nm) using a confocal microscopy system (Leica TCS SP5).

### 2.7 Immunofluorescence studies

Cultured PASMCs were fixed with 2% paraformaldehyde, permeabilized with 0.05% Triton X-100, and incubated with primary antibodies (24 h at 4°C) followed by secondary antibodies (24 h at 4°C) as indicated in Table 1. Samples were mounted with FluoroGel (Electron Microscopy Sciences) and images were acquired using confocal microscopy (Leica TCS SP5).

### 2.8 Mitochondrial bioenergetics and metabolic profile analyses

Oxygen consumption rate (OCR) and extracellular acidification rate (ECAR) were measured in PASMCs from wild-type (*Asic1a⁺/⁺*) and *Asic1a* knockout (*Asic1a⁻/⁻*) mice using a Seahorse XF Cell Mito Stress Test Kit (Agilent), as previously described (23). For *in vitro* hypoxia, PASMCs were transiently cultured in a hypoxic subchamber (BioSpherix) at 2% O₂, 5% CO₂, and balance N₂ for 72 h.

Following calibration, OCR and ECAR were measured simultaneously. After basal OCR was measured, samples were treated with 1 μM oligomycin (Table 2) to determine ATP production. Maximal respiration was determined following the addition of 1 μM FCCP, and the spare respiratory capacity is calculated as the difference between maximal respiration and basal respiration. Injection of the complex I and III inhibitors, rotenone and antimycin A (500 nM each), respectively, enables the calculation of nonmitochondrial respiration and proton leak. Coupling efficiency was expressed as the ratio of ATP production to basal respiration (as a percentage).

**Table 2.**
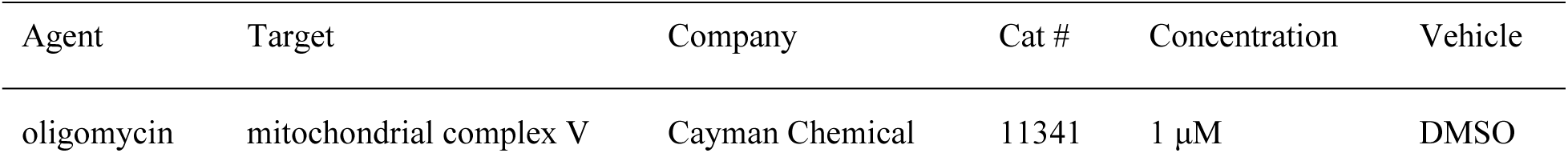
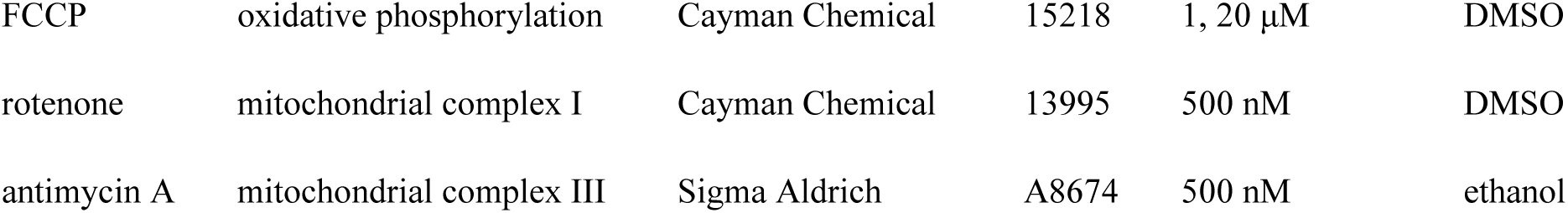
List of pharmacological agents and concentrations used.

| Agent | Target | Company | Cat # | Concentration | Vehicle |
| --- | --- | --- | --- | --- | --- |
| oligomycin | mitochondrial complex V | Cayman Chemical | 11341 | 1 μM | DMSO |
| FCCP | oxidative phosphorylation | Cayman Chemical | 15218 | 1, 20 μM | DMSO |
| rotenone | mitochondrial complex I | Cayman Chemical | 13995 | 500 nM | DMSO |
| antimycin A | mitochondrial complex III | Sigma Aldrich | A8674 | 500 nM | ethanol |

Additionally, parallel experiments were performed using a Seahorse XF Cell Energy Phenotype Test (Agilent). Basal OCR and ECAR were measured to establish baseline metabolic phenotype, followed by oligomycin and FCCP treatment to induce a stressed phenotype. The metabolic potential was expressed as the ratio of stressed to basal values.

### 2.9 Mitochondrial Morphology

IPAs were pressurized to 15 mm Hg, as we previously described (13), and fixed in 3% formaldehyde and 2% glutaraldehyde (electron microscopy grade) in 0.1 M sodium phosphate buffer (pH 7.4), washed, and post-fixed for 1 h in 1% osmium tetroxide and 0.1 M imidazole (pH 7.4). Samples were rinsed in buffer and ultrapure water, then dehydrated through a graded ethanol series. Samples were subsequently transitioned to propylene oxide and embedded in Epon-Araldite resin. Thin sections (60–80 nm) were imaged using a Hitachi HT7700 transmission electron microscope equipped with an AMT XR16M 16-megapixel camera. Mitochondria within PASMCs were identified, and mitochondrial morphometric analysis was performed in ImageJ using the polygon selection tool.

### 2.10 Calculations and statistical analysis

All data are expressed as means ± SEM. Statistical comparisons were made using Prism 10 (GraphPad Software). The statistical tests used and exact p-values are reported in the figure legends. A probability of <0.05 with a power level of 0.80 was accepted as statistically significant for all comparisons.

## 3 Results

### 3.1 Chronic hypoxia induces mitochondrial dysfunction in PASMCs

We first examined the effects of CH on mitochondrial function in PASMCs. TMRE fluorescence, an indicator of ΔΨm, was significantly elevated in PASMCs from CH rats compared with normobaric controls, consistent with ΔΨm hyperpolarization (Figure 1A–B). This was accompanied by increased Rhod-2 AM fluorescence, indicating higher mitochondrial Ca²⁺ levels (Figure 1C–D), and increased MitoSOX fluorescence, reflecting greater mitochondrial O_2_^-^ generation (Figure 1E–F). In contrast, cleaved caspase-3 immunofluorescence was reduced in PASMCs from CH rats, suggesting impaired activation of the intrinsic apoptotic pathway (Figure 1G–H). Together, these findings indicate that CH promotes ΔΨm hyperpolarization, elevated [Ca²⁺]*_mito_*, and oxidative stress in PASMC mitochondria while suppressing caspase activation.

**Figure 1.**
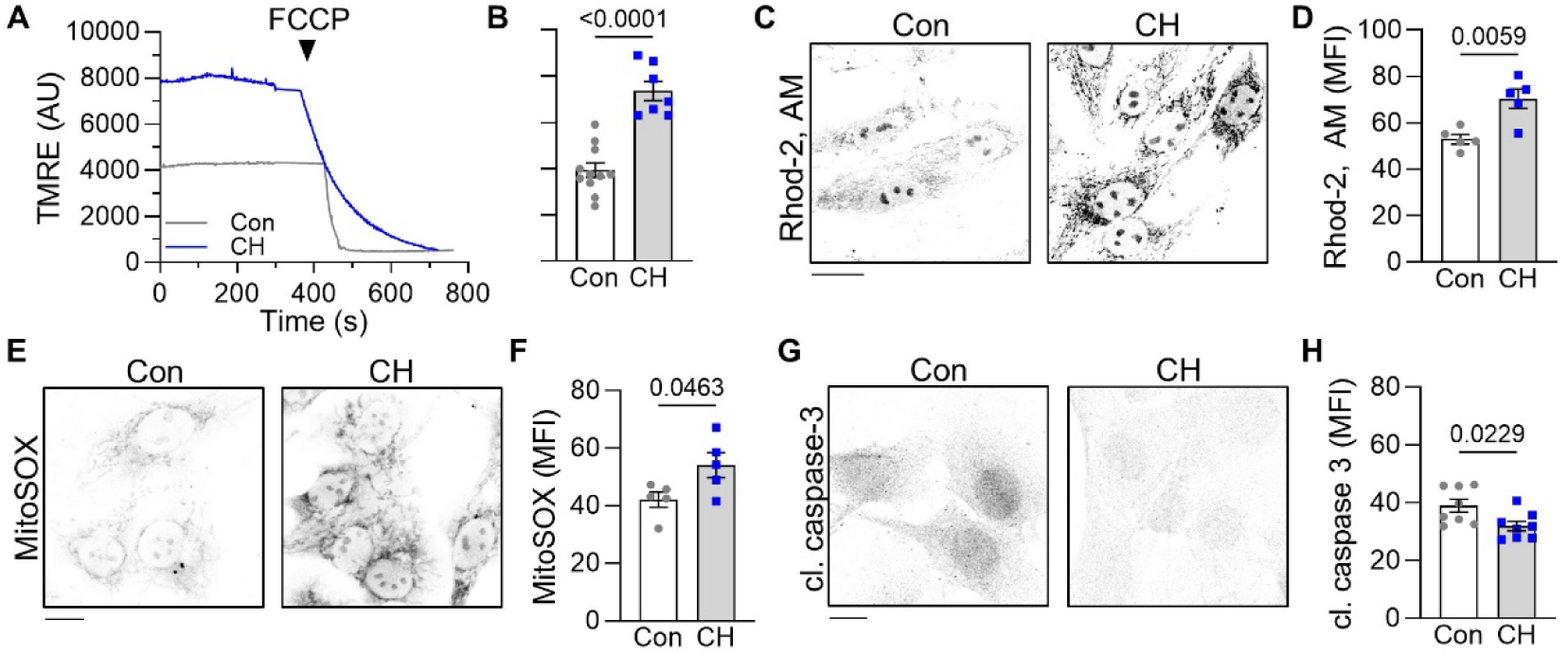
CH causes mitochondrial impairment and inhibits caspase cleavage. A) Representative traces and B) summary data for TMRE fluorescence in PASMCs from control (Con, gray circles) and CH (blue squares) rats. Following baseline recordings, 20 μM FCCP was superfused to uncouple oxidative phosphorylation and collapse ΔΨm. Rhod-2, AM and MitoSOX Red were used to determine relative mitochondrial Ca^2+^ (C and D) and mitochondrial O2-(E and F) levels, respectively. G) Representative images and H) summary data showing protein expression of cleaved caspase-3 by immunofluorescence. Values are means ± SEM; data points indicate *n* as the number of animals/group; analyzed by unpaired t test; image scale bars=10 μm.

To determine whether these mitochondrial disturbances were associated with altered mitochondrial quality control, we assessed markers of mitophagy and mitochondrial dynamics. The microtubule-associated protein 1 light chain 3 beta (LC3B)-II/I ratio and mitochondrial Parkin levels were both reduced in IPAs from CH rats, consistent with impaired mitophagy (Figure 2A–C). Western blot analysis of mitochondrial fusion proteins revealed increased mitofusion 1 (MFN1) expression, whereas mitofusion 2 (MFN2) was unchanged. Moreover, expression of the mitochondrial fission proteins dynamin-related protein 1(DRP1) and Fission 1 (FIS1) was unchanged (Figure 2D–E). Transmission electron microscopy of PASMC mitochondria from pressurized IPAs demonstrated enlarged mitochondria with reduced number per cell in CH rats, as reflected by increased area and circularity, and decreased aspect ratio (Figure 2F–G). These results suggest that CH disrupts mitochondrial morphology and quality control, potentially leading to an accumulation of dysfunctional mitochondria.

**Figure 2.**
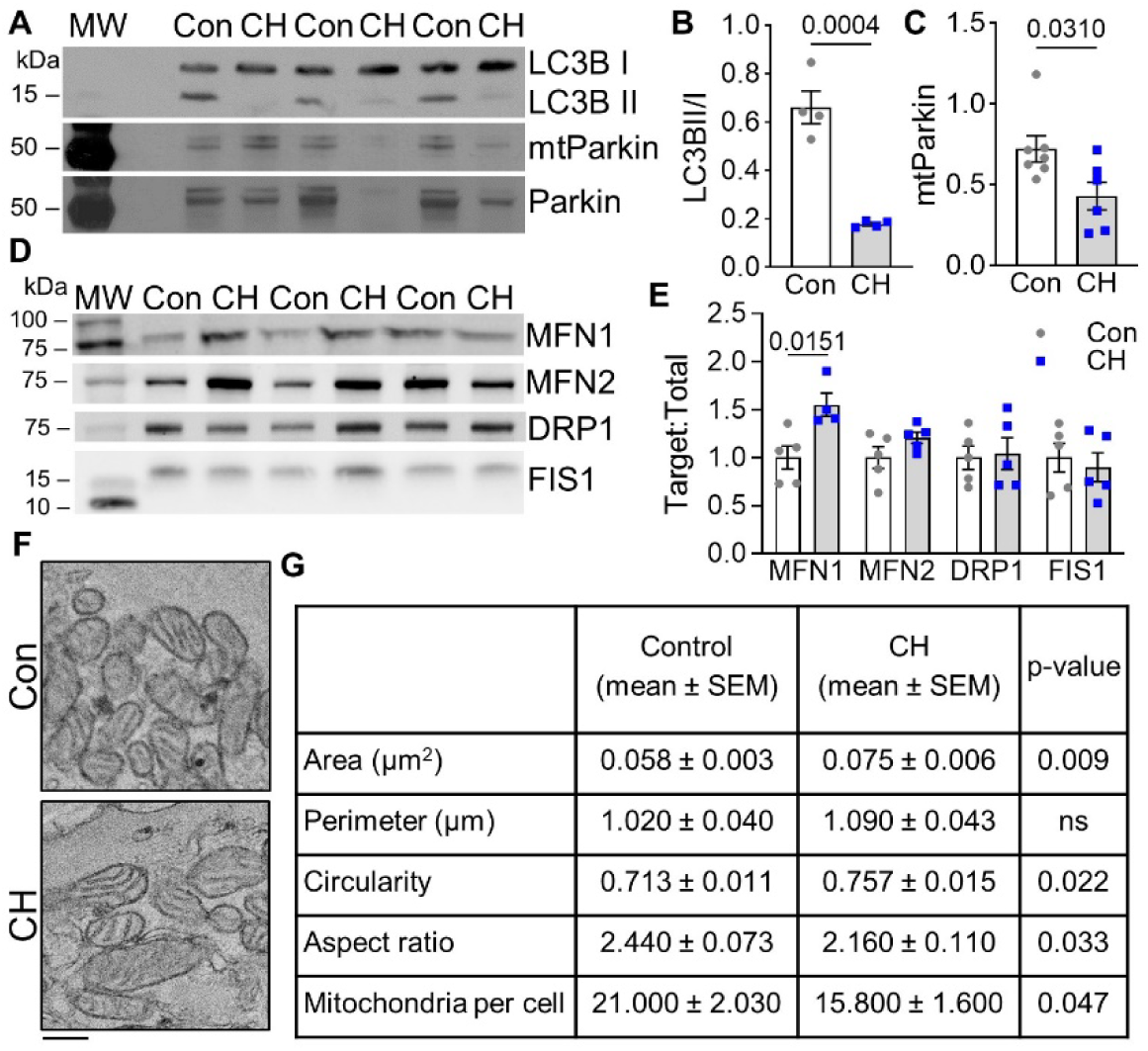
CH inhibits mitochondrial quality control. A) Representative western blots and summary data of B) the ratio between LC3B II (conjugated-form; ∼14kDa) and LC3B I (unconjugated-form; ∼17 kDa) and C) mitochondrial Parkin (∼52 kDa) normalized to cellular Parkin levels in PASMCs from control (Con, gray circles) and CH (blue squares) rats. D) Representative western blots and E) summary data showing protein expression of MFN1 (∼84 kDa), MFN2 (∼86 kDa), DRP1 (∼82 kDa), and FIS1 (∼17 kDa) in IPAs from control and CH rats. Protein expression was normalized to total lane protein determined by CBB staining. F) Representative transmission electron microscope images and G) summary data of mitochondrial morphology and numbers in PASMCs within pressurized IPAs from control and CH rats. Values are means ± SEM; data points indicate *n* as the number of animals/group; analyzed by unpaired t test; image scale bars=0.2 μm.

### 3.2 Chronic hypoxia depletes mitochondrial ASIC1a

Consistent with prior reports in neurons (19), we show ASIC1a is expressed in mitochondrial fractions from PASMCs and IPAs. Importantly, there are markedly lower mtASIC1a protein levels in CH rats compared with controls (Figure 3A–B). Immunofluorescence microscopy confirmed this finding, showing that CH decreased the co-localization of ASIC1a with the mitochondrial marker TOMM20 in PASMCs (Figure 3C–D). These data suggest that CH selectively depletes mtASIC1a without changing total cellular expression (18).

**Figure 3.**
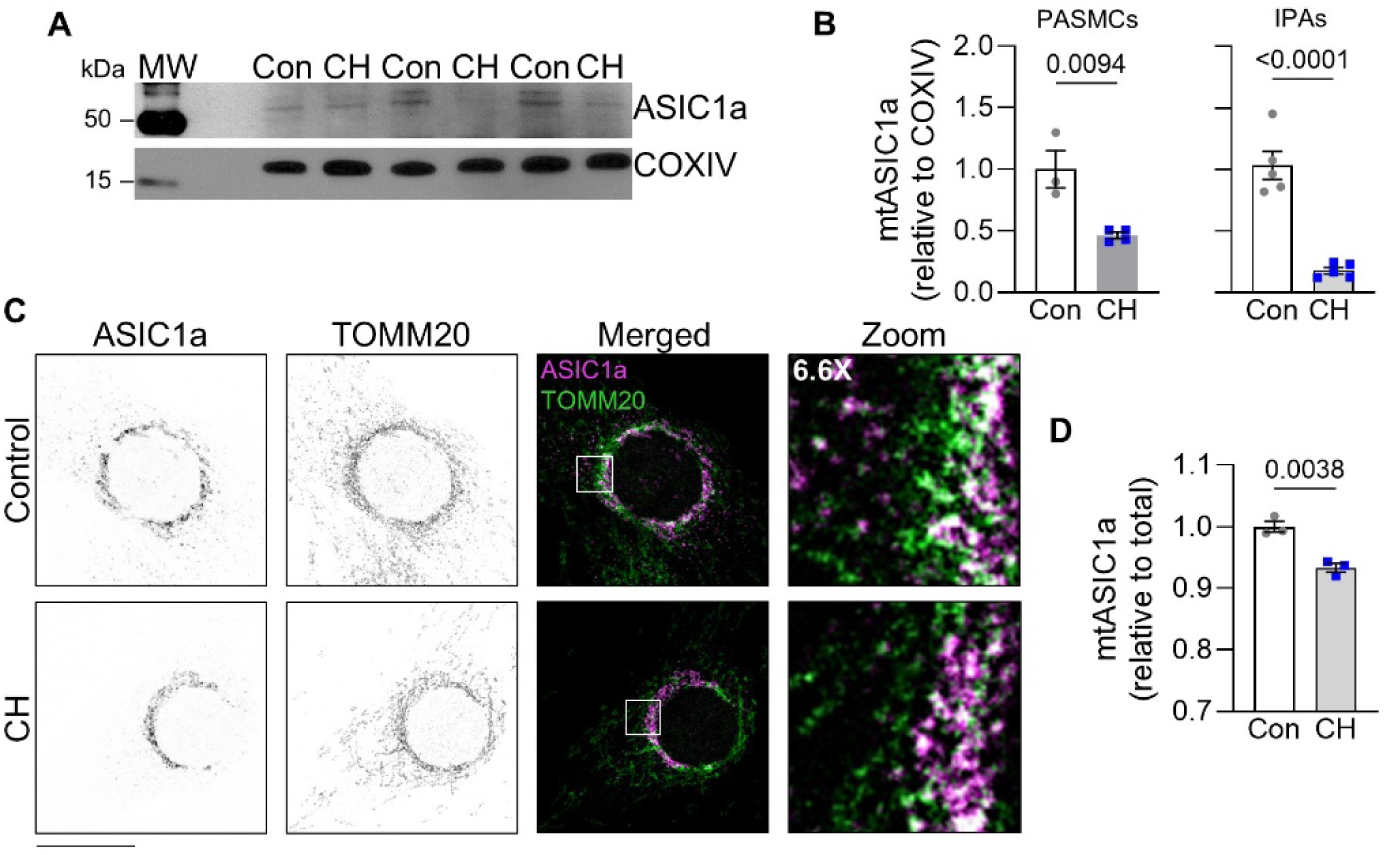
CH decreases mtASIC1a localization. A) Representative western blot and B) summary data showing protein expression of mitochondrial ASIC1a normalized to cytochrome c oxidase subunit IV (COXIV) levels in PASMCs (left) and IPAs (right) from control (Con, gray circles) and CH (blue squares) rats. C) Representative images and D) summary data for relative mtASIC1a levels determined by immunofluorescence and colocalization (indicated in white) of ASIC1a and TOMM20 in PASMCs from control and CH rats. Regions of interest were created using TOMM20 immunofluorescence. ASIC1a immunofluorescence was calculated and expressed as the ratio between ROI mean intensity and total mean intensity. Values are means ± SEM; data points indicate *n* as the number of animals/group; analyzed by unpaired t test; image scale bar=20 μm.

### 3.3 Genetic deletion of ASIC1a reproduces mitochondrial dysfunction

To determine the role of ASIC1a in regulating mitochondrial function without the confounding effects of CH, we studied PASMCs from *Asic1a^+/+^* and *Asic1a^-/-^* mice. Similar to CH rats, PASMCs from *Asic1a^-/-^* mice exhibited increased TMRE (Figure 4A-B), Rhod-2, AM (Figure 4C-D), and MitoSOX fluorescence (Figure 4E-F), indicating ΔΨm hyperpolarization, Ca²⁺ accumulation, and increased O_2_^-^ generation.

**Figure 4.**
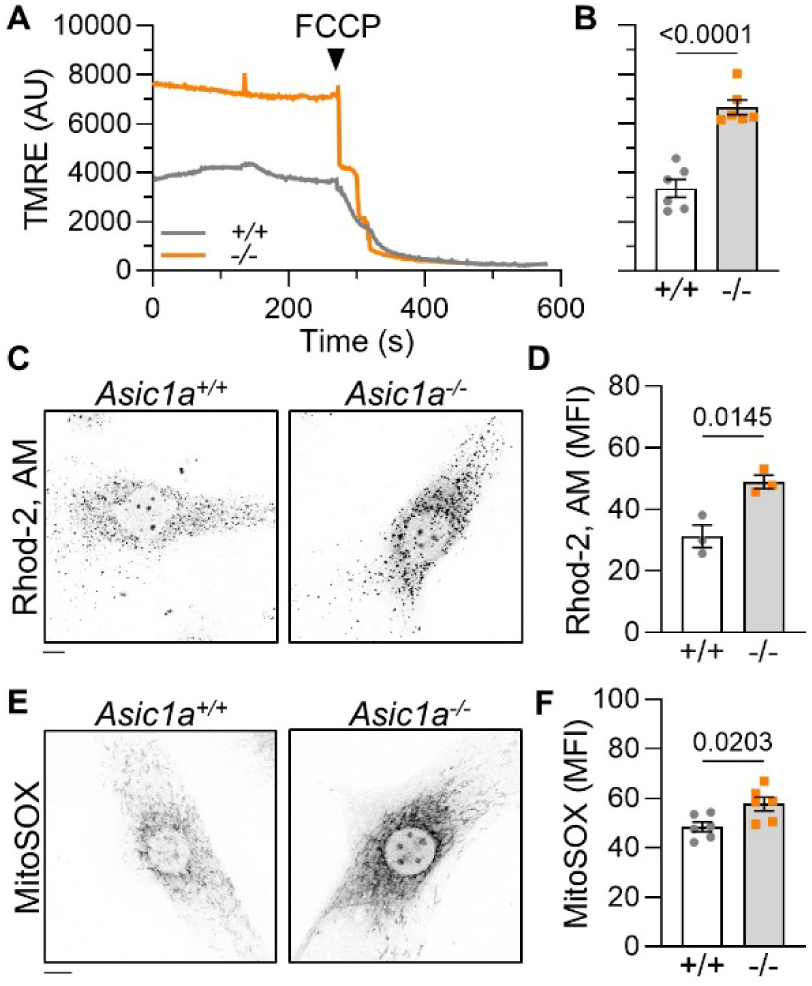
Genetic deletion of ASIC1a induces mitochondrial dysfunction. A) Representative traces and B) summary data for TMRE fluorescence in PASMCs from *Asic1a^+/+^* (+/+, gray circles) and *Asic1a^-/-^* (-/-, orange squares) mice. Following baseline recordings, 20 μM FCCP was superfused to uncouple oxidative phosphorylation and collapse ΔΨm. Representative images and summary data for C-D) Rhod-2, AM and E-F) MitoSOX fluorescence. Values are means ± SEM; data points indicate *n* as the number of animals/group; analyzed by unpaired t test; image scale bars=10 μm.

Seahorse bioenergetics analysis revealed further alterations in mitochondrial function. PASMCs from *Asic1a^-/-^* mice displayed elevated basal respiration and ATP production under both normoxia and following *in vitro* hypoxia (2% O_2_ for 72 h; Figure 5B–C). However, hypoxia independently reduced basal and maximal respiration, spare respiratory capacity, and ATP production regardless of genotype (Figure 5B–C), whereas proton leak and coupling efficiency were unchanged (Figure 5D–E). Notably, under hypoxic conditions, ECAR was elevated in *Asic1a^+/+^* PASMCs and was further increased in *Asic1a^-/-^* PASMCs, consistent with a glycolytic shift (Figure 5F). Consistent with these findings, normoxic PASMCs from *Asic1a^-/-^* mice retained a more oxidative phenotype under both basal and stressed conditions. However, *Asic1a* deletion exacerbated the shift to a glycolytic phenotype following hypoxia (Figure 5G). Together, these findings suggest that ASIC1a deficiency is accompanied by mitochondrial stress.

**Figure 5.**
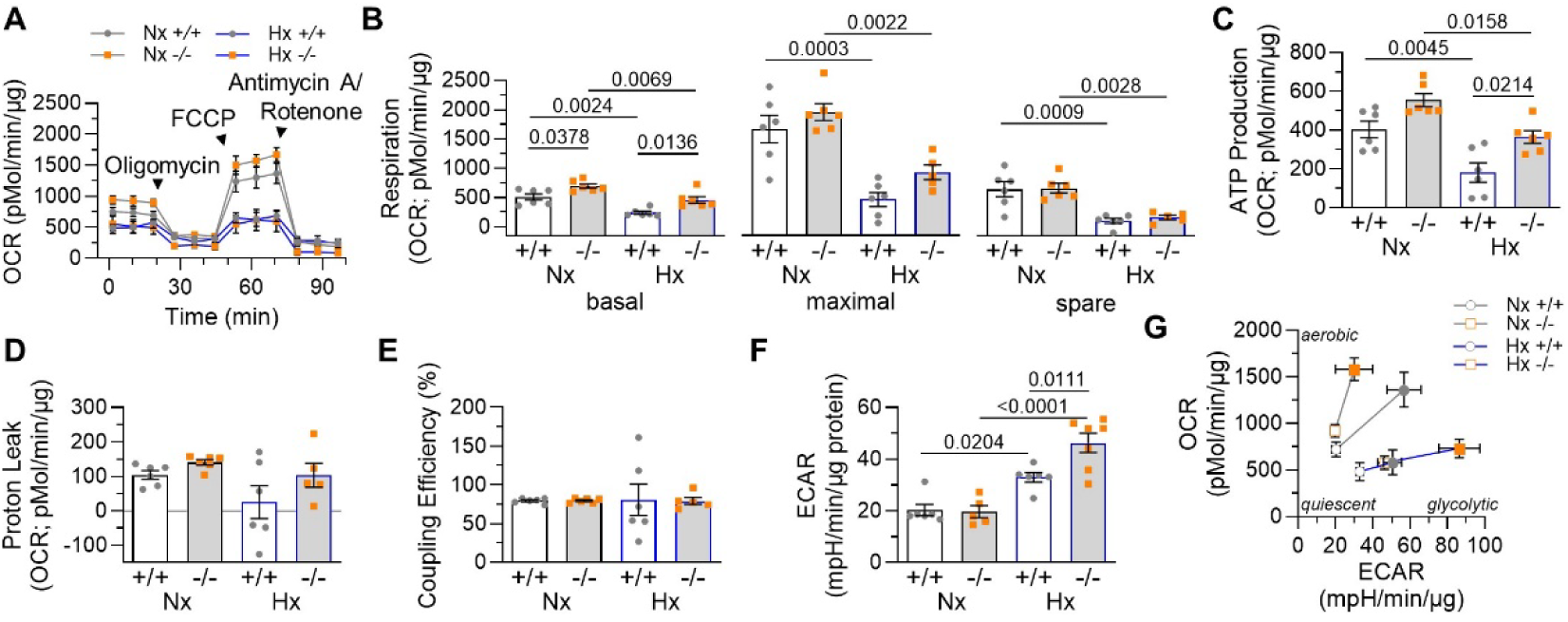
Inhibition of mitochondrial respiration following hypoxia is ASIC1a-independent. A) Seahorse XF Cell Mito Stress test was used to measure OCR (pMol/min/µg protein) over time under basal conditions and after the addition of mitochondrial inhibitors (1 µM oligomycin, 1 µM FCCP and 0.5 µM rotenone/antimycin) in PASMC from *Asic1a^+/+^* (+/+, gray circles) and *Asic1a^-/-^* (-/-, orange squares) mice. B) Assessment of basal respiration, maximal respiration, and spare respiratory capacity; C) ATP production; D) proton leak; E) mitochondrial-ATP coupling efficiency (%); and F) ECAR (mpH/min/µg protein). G) Seahorse XF Cell Energy Phenotype Test was used to measure OCR and ECAR under basal conditions (open symbols) and stressed conditions (closed symbols) in PASMCs from *Asic1a^+/+^* and *Asic1a^-/-^*mice. Values are means ± SEM; data points indicate *n* as the number of animals/group; analyzed by two-way ANOVA followed by Tukey’s multiple comparisons test.

In parallel, IPAs from *Asic1a^-/-^* mice displayed reduced LC3BII/I but increased PINK1 levels, further supporting impaired mitophagy (Figure 6A–C). Although expression of mitochondrial dynamics proteins (MFN1, MFN2, DRP1, FIS1) was unchanged (Figure 6D–E), electron microscopy confirmed mitochondrial abnormalities resembling those in CH rats, including increased size and circularity, decreased aspect ratio, and reduced mitochondrial number per cell (Figure 6F–G). Thus, *Asic1a* deletion phenocopies the mitochondrial structural and quality control defects induced by CH.

**Figure 6.**
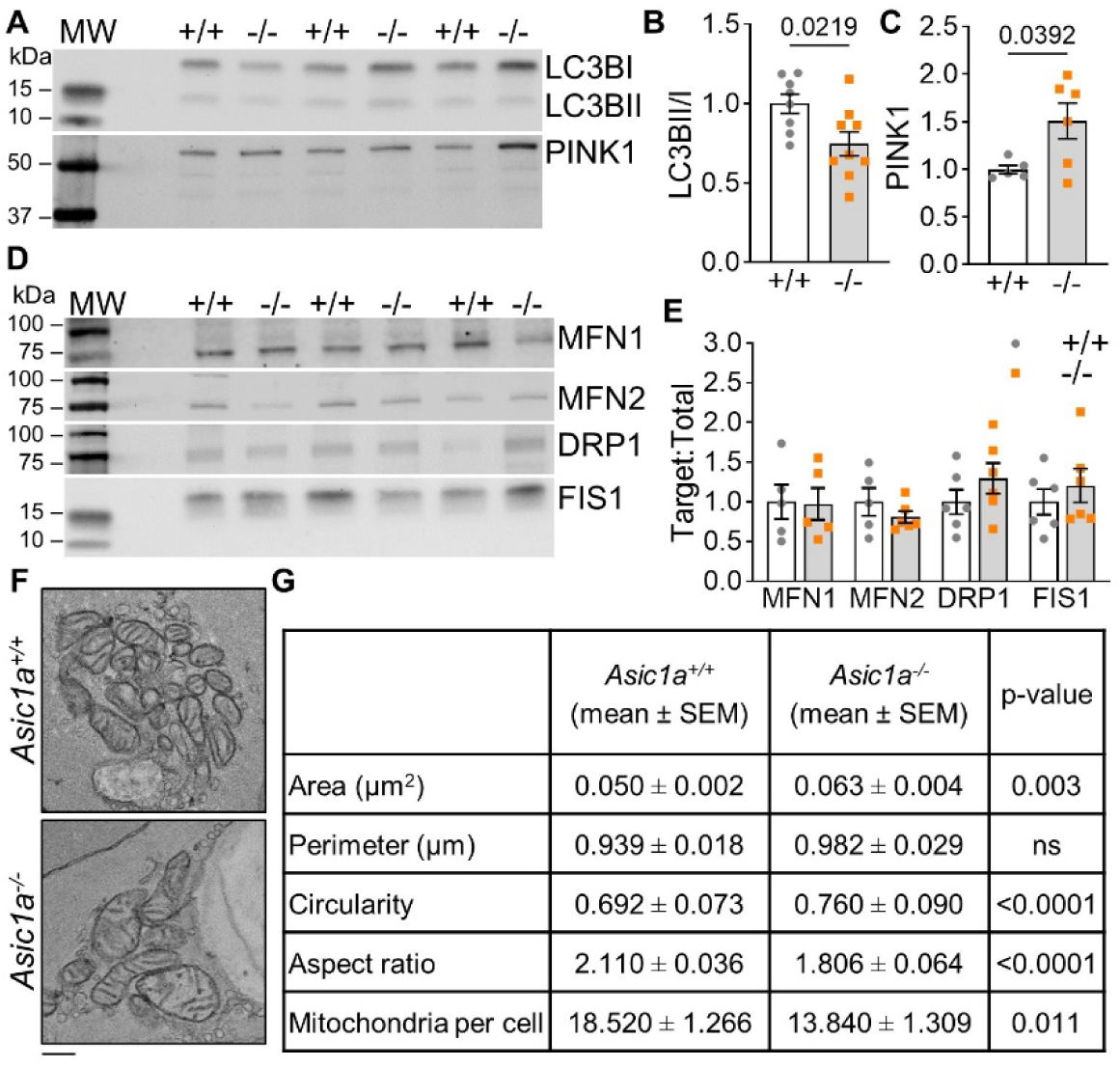
ASIC1a deficiency inhibits mitophagy and alters morphology. A) Representative western blots and summary data of B) LC3B II relative to LC3B I, C) PINK1 (∼63 kDa), and D-E) MFN1, MFN2, DRP1, and FIS1 expression in IPAs from *Asic1a^+/+^* (+/+, gray circles) and *Asic1a^-/-^* (-/-, orange squares) mice. Protein expression was normalized to total lane protein determined by CBB staining. F) Representative transmission electron microscope images and G) summary data of mitochondria morphology and numbers in PASMCs within pressurized IPAs from *Asic1a^+/+^* and *Asic1a^-/-^* mice. Values are means ± SEM; data points indicate *n* as the number of animals/group; analyzed by unpaired t test; image scale bars=0.2 μm.

### 3.4 Mitochondria-targeted ASIC1a rescues mitochondrial dysfunction

To specifically restore mtASIC1a, we transduced PASMCs with EGFP-mtASIC1a. Following validation studies that showed increased mtASIC1a localization (Figure 7A–B), we sought to determine the effects on mitochondrial function. We found that ΔΨm and caspase-3 cleavage were restored to control levels in *Asic1a^-/-^* PASMCs transduced with EGFP-ASIC1a compared with EGFP-empty (Figure 8A–D).

**Figure 7.**
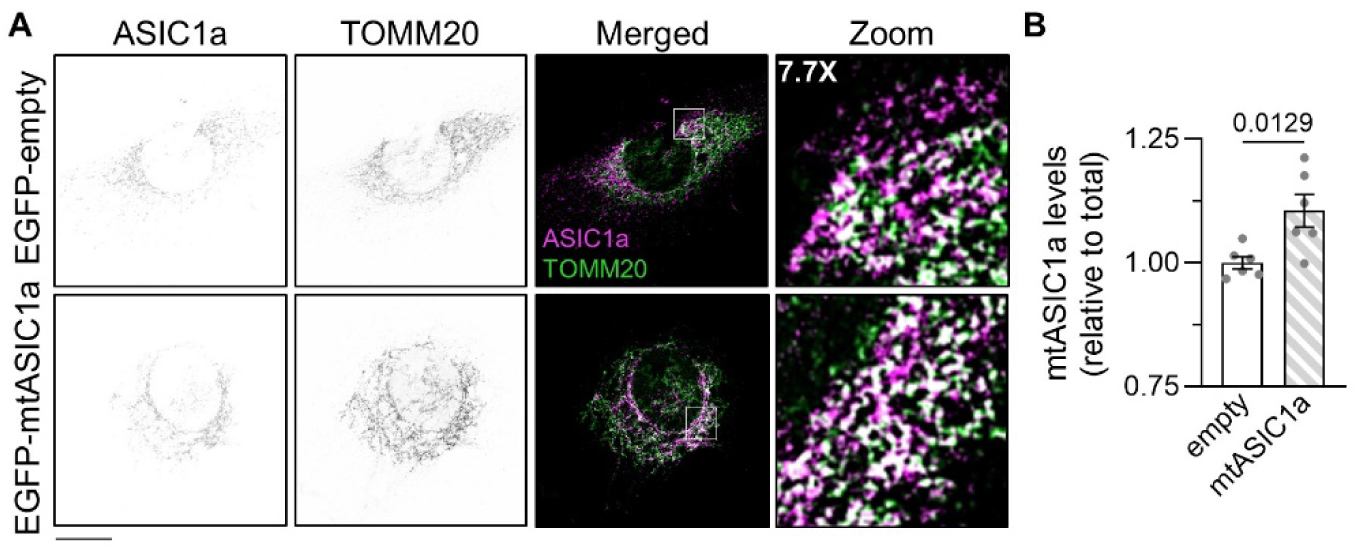
mtASIC1a localization is increased in PASMCs via lentiviral transduction. A) Representative images and B) summary data for relative mtASIC1a levels determined by immunofluorescence and colocalization (indicated in white) of ASIC1a and TOMM20 in PASMCs transduced with EGFP-empty (solid bars) or EGFP-mtASIC1a (striped bars). ROI were created using TOMM20 immunofluorescence images. ASIC1a immunofluorescence was calculated and expressed as the ratio between ROI mean intensity and total mean intensity. Values are means ± SEM; data points indicate *n* as the number of animals/group; analyzed by unpaired t test; image scale bar=20 μm.

**Figure 8.**
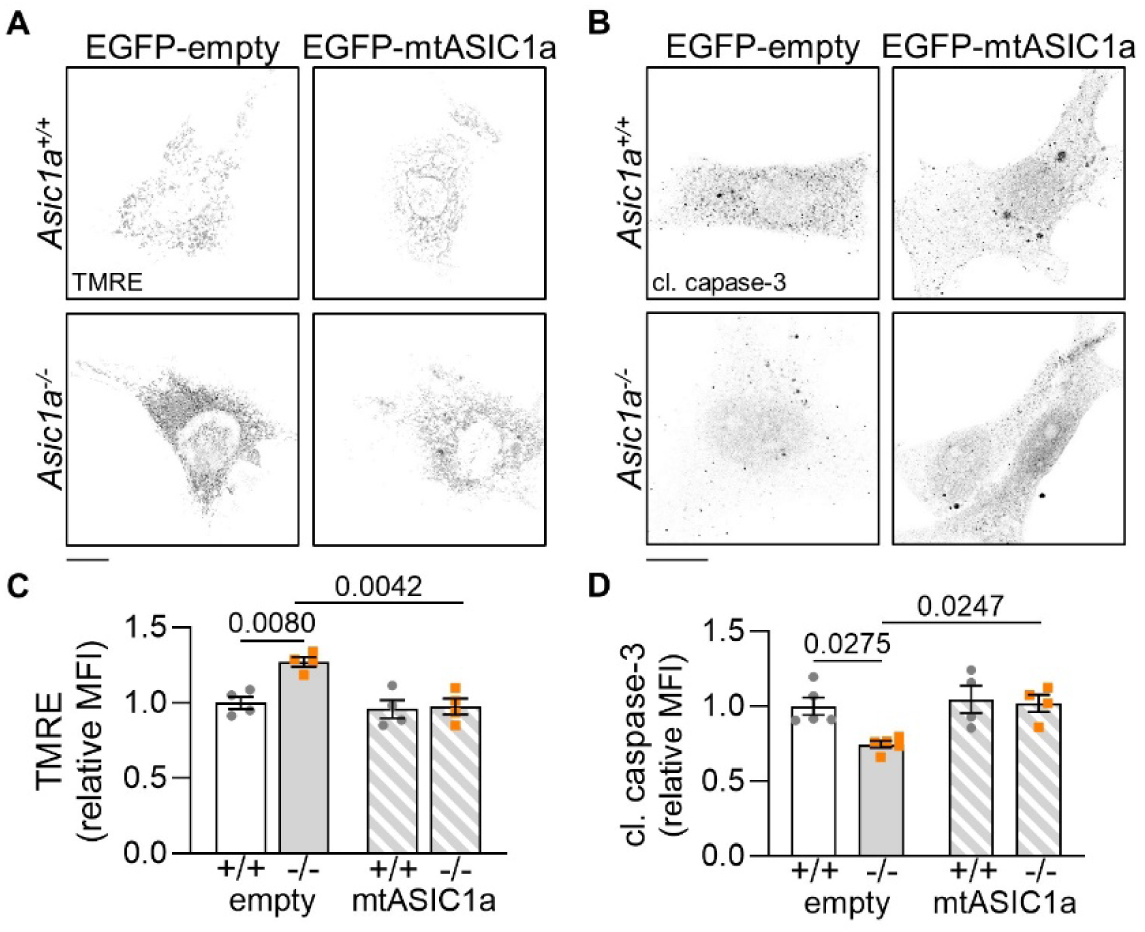
EGFP-mtASIC1a normalizes mitochondrial function in ASIC1a-deficient PASMCs. Representative images and summary data for A and C) TMRE fluorescence and B and D) cleaved caspase-3 immunofluoresence in PASMCs transduced with EGFP-empty (solid bars) or EGFP-mtASIC1a (striped bars) from *Asic1a^+/+^*(+/+, gray circles) and *Asic1a^-/-^* (-/-, orange squares) mice. Values are means ± SEM; data points indicate *n* as the number of animals/group; analyzed by two-way ANOVA followed by Tukey’s multiple comparisons test; image scale bar=20 μm.

Next, we evaluated whether EGFP-mtASIC1a could rescue mitochondrial dysfunction following CH. Significantly, ΔΨm hyperpolarization was prevented in PASMCs from CH rats by EGFP-mtASIC1a (Figure 9A and C). Moreover, pro-apoptotic signaling was restored, as evidenced by increased immunofluorescence of cleaved caspase-3 (Figure 9B and D). Together, these findings demonstrate that mtASIC1a is both necessary and sufficient to maintain mitochondrial homeostasis and promotes caspase activity involved in apoptotic signaling in PASMCs.

**Figure 9.**
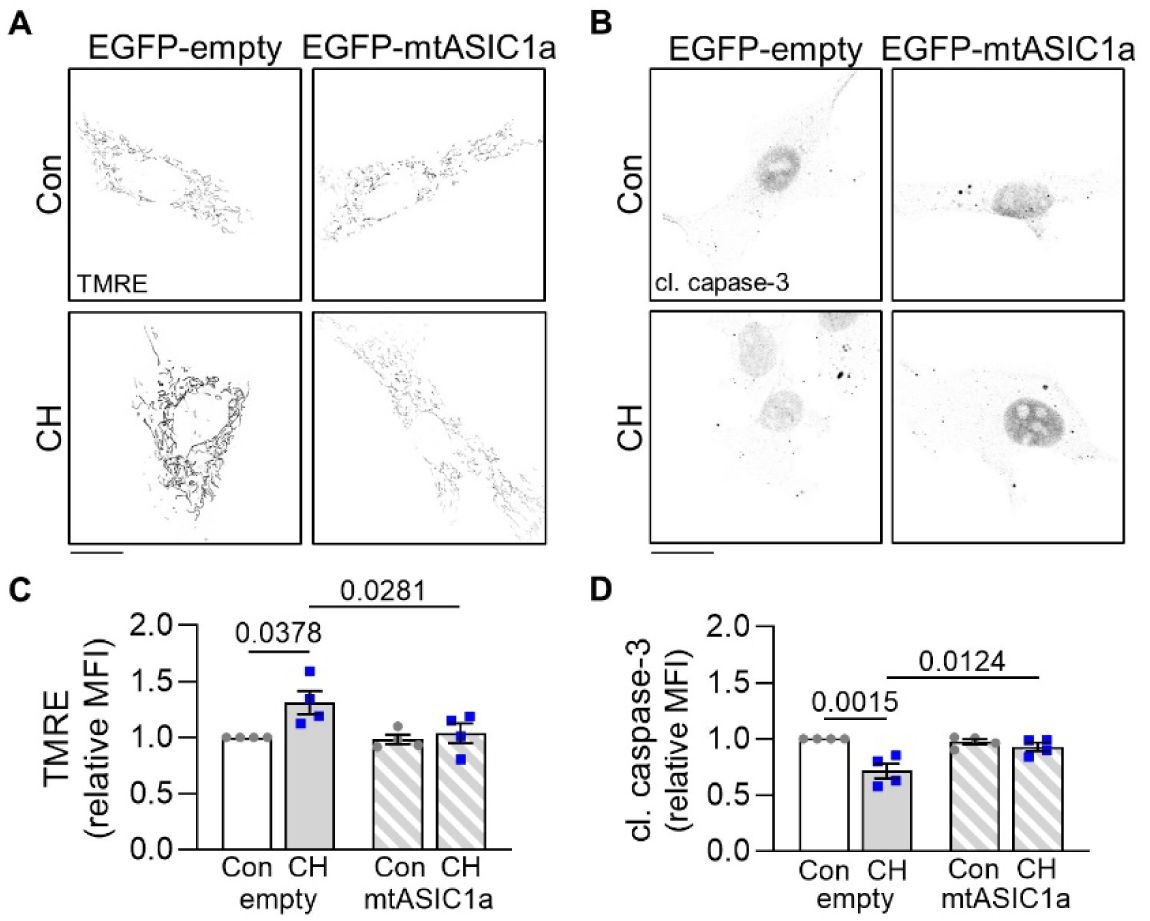
EGFP-mtASIC1a rescues CH-induced mitochondrial dysfunction. Representative images and summary data for A and C) TMRE fluorescence and B and D) cleaved caspase-3 immunofluoresence in PASMCs transduced with EGFP-empty (solid bars) or EGFP-mtASIC1a (striped bars) from control (Con, gray circles) and CH (blue squares) rats. Values are means ± SEM; data points indicate *n* as the number of animals/group; analyzed by two-way ANOVA followed by Tukey’s multiple comparisons test; image scale bar=20 μm.

## 4 Discussion

Mitochondrial dysfunction is increasingly recognized as a central driver of PH pathogenesis. Functional and structural remodeling of mitochondria favors proliferation, contraction, and resistance to apoptosis in pulmonary vascular cells, while also reshaping right ventricular energetics (9–12). Mitochondrial ion channels are vital for maintaining mitochondrial and cellular function, yet they have been historically understudied, especially in PH.

Our current findings identify the non-selective cation channel ASIC1a as a mitochondrial ion channel that regulates mitochondrial homeostasis in PASMCs. Specifically, we found that mtASIC1a depletion, either by CH or *Asic1a* deletion, is associated with ΔΨm hyperpolarization and subsequent mitochondrial dysfunction. Loss of mtASIC1a was further linked with mitochondrial calcium and reactive oxygen species (ROS) accumulation, impaired mitophagy, and inhibition of caspase signaling, all features that align with apoptosis resistance and pro-survival pathways central to vascular remodeling. Significantly, restoring mtASIC1a via lentiviral transduction reversed ΔΨm hyperpolarization and rescued cleaved caspase-3 levels, underscoring its functional significance in maintaining mitochondrial signaling pathways that regulate cell fate.

Mitochondrial ion homeostasis is tightly regulated by a complex interplay of outer and inner membrane channels and transporters; however, the functional contribution of specific channels remains poorly understood. Notably, the identification of putative mitochondrial proteins and their definitive roles in PH pathogenesis remains challenging. As such, studies have historically focused on channels/transporters present exclusively in mitochondria, including voltage-dependent anion channels (VDAC), the mitochondrial calcium uniporter (MCU) complex, and uncoupling proteins (UCP) (24–26). Interestingly, some studies suggest that ATP-sensitive K^+^ channels, Cl^-^ intracellular channel 1 and −4, and the Ca^2+^-activated Cl^-^ channel, Anoctamin-1, localize to mitochondria and participate in pulmonary vascular cell homeostasis (27–29). Along with VDAC, MCU, and UCP, their coordinated activities influence several key mitochondrial processes, including energy metabolism, ΔΨm, Ca^2+^ homeostasis, and redox balance. Our findings suggest that mtASIC1a participates in mitochondrial cation conductance, as its depletion consistently produced ΔΨm hyperpolarization. This is consistent with the notion that, as a component of the inner mitochondrial membrane (19), ASIC1a, along with other ion channels, elicits mild depolarizing currents in response to the highly negative electrochemical potential. As such, impaired cation influx or anion efflux exaggerates ΔΨm, potentially predisposing mitochondria to Ca^2+^ accumulation and oxidative stress.

Many PH models exhibit ΔΨm hyperpolarization in PASMCs, and sometimes in the endothelium (30,31). This highly negative state favors increased Ca^2+^ uptake, predominantly by the MCU complex, and is directly involved in dehydrogenase activation and ROS generation (32,33). Our findings that [Ca^2+^]*_mito_* is increased in PASMCs from CH rats and *Asic1a^-/-^* mice parallel previous work showing that Ca^2+^-induced Ca^2+^ uptake is increased in neuronal mitochondria from *Asic1a^-/-^* mice, which may suggest enhanced conductance and/or increased Ca^2+^ retention capacity (19). Additionally, we found increases in [O_2_^-^]*_mito_* following CH or *Asic1a* deletion, but the modality may vary between experimental conditions. *Asic1a* deletion augmented mitochondrial respiration and ATP production, which may stimulate electron flow and ROS generation; however, under hypoxic conditions, impaired mitochondrial respiration persisted regardless of genotype. These findings suggest that the metabolic shift in PASMCs following hypoxia occurs independently of ASIC1a. Specifically, stabilization of hypoxia-inducible factor increases pyruvate dehydrogenase kinase, leading to the inhibition of pyruvate conversion. As such, resultant impairment of the citric acid cycle reduces mitochondrial-dependent glucose utilization, which is accompanied by increased glycolysis and lactic acid fermentation (34). Importantly, we previously showed this metabolic shift leads to extracellular acidification and potentiates ASIC1a activity at the plasma membrane (23).

Depending on the duration and severity, elevated [Ca^2+^]*_mito_* and ROS can initiate either pro-apoptotic or pro-survival cellular responses (35,36). Significantly, these factors can promote mitochondrial permeability transition pore (mPTP) opening, which is essential for cytochrome C mobilization and caspase activation in the intrinsic apoptotic pathway. However, several studies demonstrate paradoxical inhibition of mPTP in PH (25,30). Alternatively, ΔΨm can also regulate mPTP activity, with ΔΨm depolarization inducing pore opening, and ΔΨm hyperpolarization decreasing the likelihood of pore opening (37,38). Our findings that cleaved caspase-3 levels are reduced following CH or *Asic1a* deletion align with previous reports showing genetic inhibition of ASIC1a attenuates H_2_O_2_-induced ΔΨm depolarization and subsequent mPTP opening, leading to decreased cell death in neurons (19). Interestingly, ASIC1a has also been implicated in acidosis-induced cell death associated with intervertebral disc degeneration, renal ischemia/reperfusion injury, and acute lung injury (39–42). However, the mechanisms underlying ASIC1a-dependent apoptosis may vary depending on the stimulus and the specific cellular pool of ASIC1a that is activated. For example, Wang et al. showed that, unlike genetic deletion, pharmacological inhibition of ASIC1a with psalmotoxin 1 did not prevent H_2_O_2_-induced cell death, despite effectively inhibiting extracellular acidosis-induced cell death (19). Since psalmotoxin 1 is a large peptide that does not readily cross the plasma membrane, these findings suggest that H_2_O_2_-induced cell death may involve intracellular ASIC1a pools rather than plasma membrane-localized channels.

Mitochondrial derangements can be exacerbated by impaired mitochondrial quality control (43,44). Our findings suggest that mtASIC1a contributes to some of these processes, including mitophagy. Other than increased MFN1 protein levels following CH, we found that fusion/fission machinery was largely unaltered, while mitophagy was impaired. Additionally, mitochondria in PASMCs from CH rats or *Asic1a^-/-^* mice show similar *in situ* morphological defects, including larger size with decreased aspect ratio and number, possibly indicating mitochondrial enlargement. While steady-state mitochondrial dynamics can significantly affect structural integrity, mitochondrial ion channel activity modulates osmotic pressure equilibrium and, therefore, is essential for maintaining mitochondrial volume homeostasis (45). Notably, increased cation influx, such as K^+^ and Ca^2+^, can increase mitochondrial matrix volume (46,47). Although understudied in the context of the pulmonary vasculature, several reports suggest that mitochondrial volume is closely related to cellular respiration, ROS formation, mitophagy, and apoptosis (45,47–50).

While our findings support the broader view that mitochondrial quality control is disrupted in PH, there are reported differences in the nature of these impairments. Our findings align with recent studies showing increased fusion and impaired mitophagy in PH (51–53). However, other studies have reported impaired fusion accompanied by increased fission, mitochondrial fragmentation, and elevated mitophagy in various models of PH (54–59). Moreover, selectively targeting these processes often improved PAEC/PASMC function; however, there was only marginal improvement in pulmonary arterial and/or right ventricular pressures in PH animals.

While these differences in mitochondrial quality control and morphology are unclear, they may stem from methodological differences, including culturing of cells, the severity, duration, and type of hypoxia (e.g., normobaric versus hypobaric, *in vivo* versus *in vitro*), and the mode of PH induction. Nonetheless, imbalances in mitochondrial quality control are increasingly recognized as a feature of PH, and our data implicate ASIC1a in these processes.

Our current findings identify mtASIC1a as a potential candidate conferring mitochondrial dysfunction and apoptosis resistance in PH. However, it is essential to note that we did not directly assess its ion conductance properties within mitochondria, which remains difficult due to limitations in inner mitochondrial membrane electrophysiological characterization. While studies in isolated mitochondria could clarify whether mtASIC1a directly mediates Na^+^/Ca^2+^ transport or modulates ΔΨm via indirect mechanisms, the goals of these studies should be carefully considered. Isolating mitochondria is technically challenging, and it is difficult to retrieve purified populations that are also functionally intact. As such, false-positive results often complicate the identification of mitochondrial ion channels. Notably, however, a recent study successfully identified ASIC1a as one of a few ion channels present in the ultrapure mitochondria fraction (20). This finding strengthens the rationale for further investigation into the direct role of ASIC1a in mitochondrial ion homeostasis and its contribution to various pathologies.

Our previous work demonstrated that ASIC1a plays an essential role in the development of CH-induced PH. Specifically, increased localization of ASIC1a to the plasma membrane promotes membrane depolarization and enhances Ca^2+^ entry, which drives PASMC proliferation and migration (13–17). The current findings build on these observations by revealing a dual, and potentially opposing, role for ASIC1a depending on its subcellular localization. Loss of mtASIC1a following CH leads to mitochondrial dysfunction and apoptosis resistance, raising the intriguing possibility that selectively enhancing mtASIC1a could restore mitochondrial function and maintain apoptosis susceptibility in pathologic PASMCs, potentially mitigating pulmonary vascular remodeling. However, both plasma membrane and mitochondrial ASIC1a contribute to PASMC dysfunction. These intersecting roles of this dual-localized ion channel suggest that selectively targeting ASIC1a trafficking may provide more effective therapeutic benefits. ASIC1a localization is regulated by cellular redox state, pH, and post-translational modifications (e.g., S-nitrosylation, S-glutathionylation, N-glycosylation) (60–63). Moreover, ASIC1a interacts with chaperones, scaffolding proteins, and cytoskeletal regulatory elements that guide subcellular localization. Indeed, we previously showed that RhoA activation in PASMCs following CH increases plasma membrane ASIC1a localization and activity (18). Future studies will further explore the molecular determinants of ASIC1a localization and their implications for PASMC function in PH.

## 5 Acknowledgments

The authors thank Tamara Howard, M.S. and Shristey Tamang for their involvement with the sample preparation (TH), electron image acquisition (TH), and image analysis (ST) for assessing mitochondria morphology. These data were generated in the HSC-Electron Microscopy Facility, which is supported by the University of New Mexico Health Sciences Center.

Additionally, we would like to acknowledge the University of New Mexico Health Sciences Center Undergraduate Pipeline Network program for providing additional support and training resources (to H.E. Medina).

## 8 Author Contributions

All persons designated as authors qualify for authorship, and all those who qualify for authorship are listed. MNT, JSN, TCR, and NLJ conceived and designed research; MNT, LMH, and NLJ performed experiments; MNT, LMH, HEM, and NLJ analyzed data; MNT, LMH, HEM, and NLJ interpreted results of experiments; MNT and NLJ prepared figures; MNT drafted manuscript; MNT, JSN, TCR, and NLJ edited and revised manuscript. All authors approved the final version of the manuscript and agree to be accountable for all aspects of the work in ensuring that questions related to the accuracy or integrity of any part of the work are appropriately investigated and resolved.

